# Deep learning-mediated detection of accelerated water drinking after aquaresis in V1b vasopressin receptor knockout mice

**DOI:** 10.64898/2026.08.21.746202

**Authors:** Hiroaki Kaminaga, Chortip Sajjaviriya, Morio Azuma, Yuji Kashiwakura, Fumihiro Niwa, Hiroyoshi Tsuchiya, Tsukasa Ohmori, Taka-aki Koshimizu

## Abstract

How water intake is initiated and maintained following V2 vasopressin receptor antagonism remains poorly understood. To elucidate the role of the V1b receptor in managing dehydration stress induced by V2 antagonism, we used deep learning-based computer vision to analyze drinking behavior in V1b knockout (V1bKO) and wild-type (WT) mice. While total water access and intake volume were comparable between genotypes, V1bKO mice exhibited distinct temporal dynamics. Modeling cumulative intake with the Hill equation revealed that the time required to reach 50% of maximal water access was significantly shorter in V1bKO mice than in WT mice. This accelerated drinking effectively mitigated increases in serum osmolality and body weight loss. A reduced Hill’s coefficient in V1bKO mice indicates a reduction of the rapid, cooperative-like water accumulation seen in WT mice. Furthermore, elevated basal hemoglobin levels in V1bKO mice were independent of dehydration, as confirmed via bone marrow transplant. Analysis of movement trajectories revealed that V1bKO mice exhibit a lower proportion of vertical movement (required for nozzle access) despite similar total distances traveled. Collectively, our results demonstrate that the V1b receptor critically regulates water-seeking behavior and osmotic homeostasis.

## 1 INTRODUCTION

The antidiuretic hormone, arginine vasopressin (AVP), plays a primary role in keeping water and electrolyte balance by transferring the aquaporin 2 channel to the luminal side of renal collecting duct cells [6, 40]. Compared to the strong anti-diuretic effect of V2 receptor activation [30, 44, 18, 9], participation of two V1-type vasopressin receptors, V1a and V1b subtype receptors, to water and electrolyte regulation has not been fully understood. In particular, a role for the V1b receptor in water homeostasis has been proposed but its precise contribution remains unclear. In the anterior pituitary, V1b receptor stimulates secretion of adrenocorticotropic hormone (ACTH) [30]. Basal and AVP-stimulated secretion of ACTH was decreased in mice lacking the V1b receptor (V1b knockout [KO] mice) [53], also see [31]. Under stress condition, the plasma ACTH and corticosterone concentrations were also decreased in the V1bKO mice after the forced swim test [53]. Because water and sodium are retained partly by the glucocorticoid and mineral corticoid system [8], the V1b receptors may contribute to water homeostasis through hypothalamus-pituitary-adrenal (HPA) axis activity in basal and stressed conditions [53, 51, 32, 33, 46].

In addition, renal V1b receptor has been reported to co-exist with V2 receptors and may counterbalance the antidiuretic effect triggered by V2 receptor activation [25, 49]. In the central nervous system, V1b receptor in the hypothalamus, subfornical organ (SFO) and organum vasculosum laminae terminalis (OVLT) may change the sensitivity to monitoring serum solute particles [20], leading to water and/or salt intake. However, the results reported from two studies were inconclusive on the consumed water and excreted urine volumes when wild-type (WT) and V1bKO mice were compared in a metabolic cage [11, 46]. To understand the V1b receptors’ multiple roles in water homeostasis and intake, and the possible inhibitory effect of the V1b receptor on the V2 receptors in the renal principal cells, the urine volumes between WT and V1bKO mice were evaluated under tolvaptan (V2 vasopressin receptor antagonist) administration [53].

In this study, deep neural networks (DNN) and computer vision techniques were utilized to monitor mouse water-access behavior within conventional, transparent glass metabolic cages. To our knowledge, this experimental setup represents a novel application based on previous metabolic studies [1]. DNN-based analysis of animal behavior has been applied successfully in studies of pain and sleep [27, 54, 15]. However, only a few studies used DNN methods to focus on water intake behavior in daily calves and pigs [19, 3]. One of advantages of DNN-based methods over the currently prevailing lickometer method is the broader detection of the water intake and other behaviors, including both successful licking and attempts of licking [2]. In our previous report, V1bKO mice moved less distance than WT when the mice were placed in a new open-filed [50]. In this study, mice were motivated by water diuresis and dehydration to initiate and repeat water intake in the metabolic cage as a new environment. This experimental set up clarifies whether internal metabolic demand, dehydration, could overcome tendency of less movement in a new field in V1bKO mice compared to WT mice.

## 2 METHODS

### 2.1 Animals

The WT and V1bKO mice were maintained in the genetic background of a hybrid 129/Sv and C57BL/6J strain [53]. The temperature of the animal room was controlled at 23*±*1*^◦^*C, the humidity at 55*±*10% and the light/dark cycle was 12/12 h. Male mice at the age of 8– 19 weeks were used. The Animal Care and Use Committee of the Jichi Medical University approved the animal experiments and experiments were conducted in accordance with the ARRIVE guidelines (Animal Research: Reporting of In Vivo Experiments).

### 2.2 Genotyping

Genomic DNA was isolated from the tails of WT or V1bKO mice. Briefly, the tails were digested with proteinase K overnight at 56*^◦^*C. Primer sequences used were as follows:

Neo primers: (sense) 5’-GTCCGGTGCCCTGAATGAACTGCAA-3’ and (antisense) 3’-ATTCGCCGCCAAGCTCTTCAG-5’.

V1bWT primers: (sense) 5’-TCTGGCCACAGGAGGC AACCT-3’ and (antisense) 3’-ATCTCGTGGCAGATGAGGCCA-5’.

The PCR cycles were as follows: an initial denaturation at 95*^◦^*C for 5 minutes, 30 cycles of denaturation at 94*^◦^*C for 30 second, annealing and extension at 60*^◦^*C for 30 second, followed by a final extension at 72*^◦^*C for 10 minutes. The PCR products were analyzed by agarose gel electrophoresis.

### 2.3 Analysis of water intake behaviors

Mouse behaviors in the metabolic cage (Sugiyama-Gen, Co., Ltd., Tokyo, Japan) were recorded for 3, 6 or 24 h with or without drug treatment. Tolvaptan (FUJIFILM Wako Pure Chemical Co., Tokyo, Japan) was dissolved at 15 mg/mL in DMSO and 20 mg/kg of tolvaptan was diluted to 100 µL by 70% ethanol. The drug or vehicle was administered intraperitoneally. The mice were placed in a cage without previous habituation to analyze their behavior responses to a new environment. Food and water were available *ad libitum*. From a video recording, 5 images of 720 x 480 pixels per 1 second were extracted by using ffmpeg program. We labelled a total of 4517 images, in which 2325 images showed mice with the intension of drinking water, and 2172 images showed mice away from the water bottle using the LabelImg graphical image annotation tool (https://github.com/tzutalin/labelImg). Supplementary Figure 1 shows examples of the labelling materials in the both situations. The labelled images were used to train DNN models using TensorFlow Object Detection API [24] and python program. The trained model’s accuracy was evaluated and further used to detect the drinking behavior of a new mouse group in a Miniconda (Anaconda, Inc.) virtual environment.

**Supplementary Figure 1:**
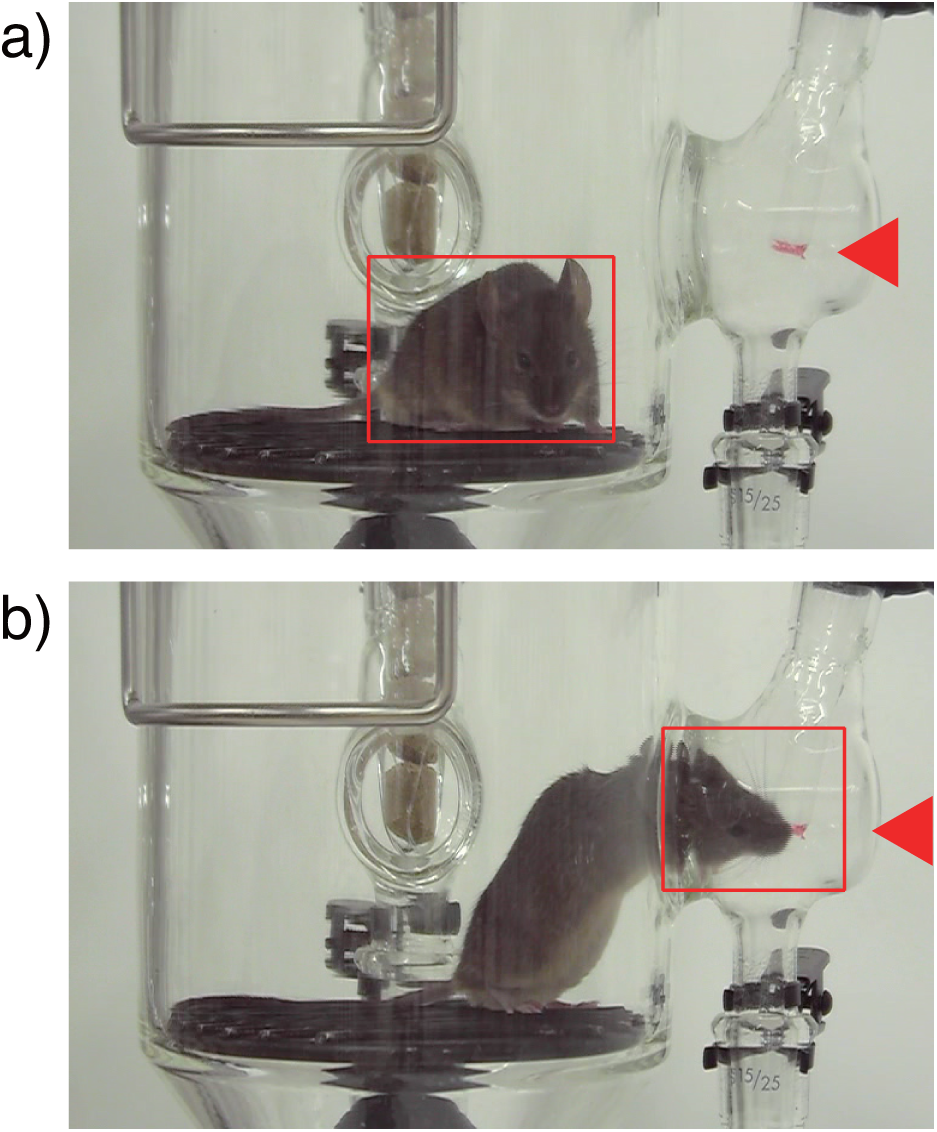
Example of object detection and classification of the mouse behaviors. Images of mouse in the metabolic cage were labelled as “away from the water” (a) or “drinking” (b). Labelled images were used for transfer learning based on Faster RCNN-Resnet101 model. Arrowheads indicate tip of the water bottle.

From the 6 h of video recording, frames were extracted at a rate of 5 fps (frames per second), resulting in about 108,000 images per mouse. The developed DNN model detected each mouse, and its behavior, either drinking or away from the water bottle, was determined. Failure of the detection was 0.2% (208 images out of 108,000 images). The rate of correct labels was 99.8%; five images were failed out of a total of 2228 images.

### 2.4 Blood chemistry and blood cell counts

Mice were deeply anesthetized with an intraperitoneal injection of pentobarbital (100 mg/kg), and blood was drawn from the inferior vena cava using a 23-gauge needle and a 1 mL syringe. Ethylenediaminetetraacetic acid dipotassium salt dihydrate (EDTA-2K) was included as anticoagulant for the measurements of white blood cell counts (WBC), red blood cell counts (RBC), haemoglobin (Hb), haematocrit (Ht), mean corpuscular volume (MCV), mean corpuscular haemoglobin (MCH), mean corpuscular haemoglobin concentration (MCHC) and platelet counts using a Celltac α MEK-6550 blood analysis system (Nihon Kohden Corp, Tokyo, Japan). Serum ion levels were measured by an EPOC^®^ blood analysis system (Siemens Healthcare Pty Ltd., Tokyo, Japan) without anticoagulant. Serum osmotic pressure was measured using a FISKE 210^®^ Osmometer (Advanced Instruments, LLC, Massachusetts, USA).

### 2.5 The enzyme-linked immunosorbent assay (ELISA) for serum erythropoietin levels

Mice were deeply anesthetized by an intraperitoneal injection of pentobarbital, and blood was drawn from the inferior vena cava. After centrifugation of the whole blood, serum was used for measurement of erythropoietin using the enzyme-linked immunosorbent assay kit (Proteintech Group, Illinois, USA), according to the manufacture’s protocol. Final absorbance at 450 nm with the correction wavelength set at 630 nm was measured by a SpectraMax M3^®^ plate reader (Molecular Devices, LLC, California, USA).

### 2.6 Bone marrow transplant (BMT)

Whole bone marrow (BM) cells were collected from WT and V1bKO mice and transplanted into irradiated WT mice, as described previously with slight modifications [41]. Briefly, the iliac, femoral and tibial bones were dipped in 70% ethanol, and BM cells were obtained in 10 mL phosphate-buffered saline (PBS). The cells were centrifuged at 2000 *×* g for 10 minutes at 4*^◦^*C. The resultant supernatant was discarded, and the collected cells were mixed in 1 mL hemolysis buffer (168 mM NH_4_Cl, 10 mM KHCO_3_, and 100 mM EDTA-Na_2_) for 5 minutes at ambient temperature. After adding 9 mL PBS, the cells were centrifuged at 2000 *×* g for 10 minutes at 4*^◦^*C. The cell pellets were mixed in 30 mL PBS, and filtrated with Cell strainer with 70 µm of mesh size (Corning Inc, Arizona, U.S.A.). After centrifugation at 2000 *×* g for 10 minutes at 4*^◦^*C, cells were resuspended at 1.0 *×* 10^7^cells/mL in PBS. For the recipient mice, 10 WT mice were irradiated with a single lethal dose of 9.5 Gy using Gamma Cell (Nordion (Canada) Inc., Ontario, Canada). For BMT, 2.0 *×* 10^6^ cells in 200 µL PBS were injected into the internal jugular vein under general anesthesia, which was initiated and maintained at 2% and 4% isoflurane, respectively. After the procedure, drink water containing 2 mg/mL neomycin and 0.05 mg/mL ampicillin was administered for 2 weeks in a pathogen-free room. Blood tests were performed 8 weeks after the BMT.

### 2.7 Measurement of circulatory plasma volume

The plasma volume of age-matched male mice was determined using the Evans blue dye dilution method, as previously described [14, 28, 48], with slight modifications. For anesthesia, a stock solution of midazolam (FUJIFILM Wako Pure Chemical Corporation) was prepared in ethanol at a concentration of 10 mg/mL. Medetomidine hydrochloride (FUJIFILM Wako Pure Chemical Corporation; 2 mg/mL) and butorphanol tartrate (Vetorphale^®^, Meiji Seika Pharma Co., Ltd., Tokyo, Japan; 5 mg/mL) were diluted in sterile saline. The anesthetic mixture was filter-sterilized and administered intraperitoneally at final doses of 2 mg/kg midazolam, 0.15 mg/kg medetomidine, and 0.5 mg/kg butorphanol. Evans blue dye (Nacalai Tesque, Inc., Kyoto, Japan; 25 µL of a 6 mg/mL solution in sterile isotonic saline) was infused into the right jugular vein using an Elite 11 syringe pump (Harvard Apparatus, Holliston, MA, USA) at a rate of 50 µL/min. Ten minutes after dye administration, blood was collected from the inferior vena cava using a heparinized syringe fitted with a 21-gauge needle and transferred to a 1.5-mL micro-centrifuge tube. Plasma was isolated by centrifugation twice at 5,000 *×* g for 5 min at 20 *^◦^*C. The resulting plasma was transferred in duplicate (100 µL per well) to a 96-well microplate, and the absorbance at 615 nm was measured using a SpectraMax M3^®^ microplate reader. Evans blue concentrations were determined from a standard curve prepared in mouse plasma, and calculated plasma volume was expressed as milliliters per gram (mL/g) of body weight.

### 2.8 Data and statistical analysis

Metabolic parameters were expressed as a box-and-whisker plot, in which the boxes indicate the 25th and 75th percentile, the middle line indicates the median, and the whiskers correspond to the minimum and maximum values. We used the R program for statistical analysis (version 4.6.1). For parametric and non-parametric comparison between the WT and V1bKO mouse groups, the Student’s t-tests and Wilcoxon tests were performed, respectively. All statistical tests were two-sided and statistical significance was defined as *p <* 0.05. The effects of genotype, condition and their interaction using a two-factor linear model (two-way ANOVA). To control for multiple testing across parameters, p-values were adjusted using the Benjamini ‒ Hochberg (BH) false discovery rate (FDR) procedure separately within each condition.

Cumulative counts of water intake time-series data without tolvaptan were grouped into uniform intervals (bins) of five seconds, expressed as the mean *±* standard error of the mean (s.e.m.), and analyzed using generalized linear mixed-effects models (GLMMs) implemented in the glmmTMB package (1.1.14) in R [7]. Because accumulating drinking counts were overdispersed relative to a Poisson distribution, a negative binomial distribution with quadratic mean-variance scaling (nbinom2) and a log link function was used for model calculations.

We treated drinking behavior over the session as non-monotonic series, and the data was fitted to models using natural cubic splines (ns(), package splines) with 2, 3, and 4 degrees of freedom, each interacted with genotype. Model fit was compared using Akaike Information Criterion (AIC) and Bayesian Information Criterion (BIC), with lower values indicating better fit relative to model complexity. The model with df = 4 provided the best fit among all candidates (AIC = 8489.1, BIC = 8618.6), a substantial improvement over the linear-time model (AIC = 8735.4, BIC = 8800.2) and over lower-order splines (df = 2: AIC = 8579.0; df = 3: AIC = 8549.1), and was retained as the final model. Statistical significance of individual fixed-effect coefficients was assessed using Wald z- tests as reported by glmmTMB, with α = 0.05.

For water access time course after tolvaptan, data without binning was fitted to Hill’s equation by Igor Pro 9 software (WaveMetrics Inc., Oregon, USA).

## 3 RESULTS

The behaviors of WT and V1bKO mice in a metabolic cage before and after administration of a tolvaptan were examined to clarify whether the V1b receptor plays a role in the retention and recovery of body water.

### 3.1 DNN analysis reveals distinct drinking behaviors in V1bKO mice compared to WT mice

From video recordings of mouse behavior, frames were extracted at a rate of 5 fps to detect mice using our trained DNN model. As detailed in the Methods section, the accuracies for detection and behavior classification were 99.8% and 99.7%, respectively.

Under baseline, freely moving conditions, V1bKO mice exhibited a trend toward reduced drinking frequency compared to WT mice, although total water access remained comparable between genotypes (Figure 1). To account for the non-independent nature of cumulative behavioral counts, we modeled discrete drinking events per interval using non-linear regression. Among models with 2, 3, and 4 degrees of freedom, the *df* = 4 model provided the optimal fit. Given the strong overdispersion observed relative to a Poisson distribution (*θ* = 0.00224), a negative binomial framework was utilized. Although no significant difference in access rate was observed at the reference time point (*z* = *−*0.217*, p* = 0.828), we identified a significant interaction between genotype and the temporal shape of water intake (genotype *×* spline term 3: *β* = *−*4.178*, SE* = 1.157, *z* = *−*3.610*, p <* 0.001). These results suggest that V1bKO mice differ from WT mice not in total water access, but in the temporal dynamics of their drinking behavior throughout the session.

**Figure 1:**
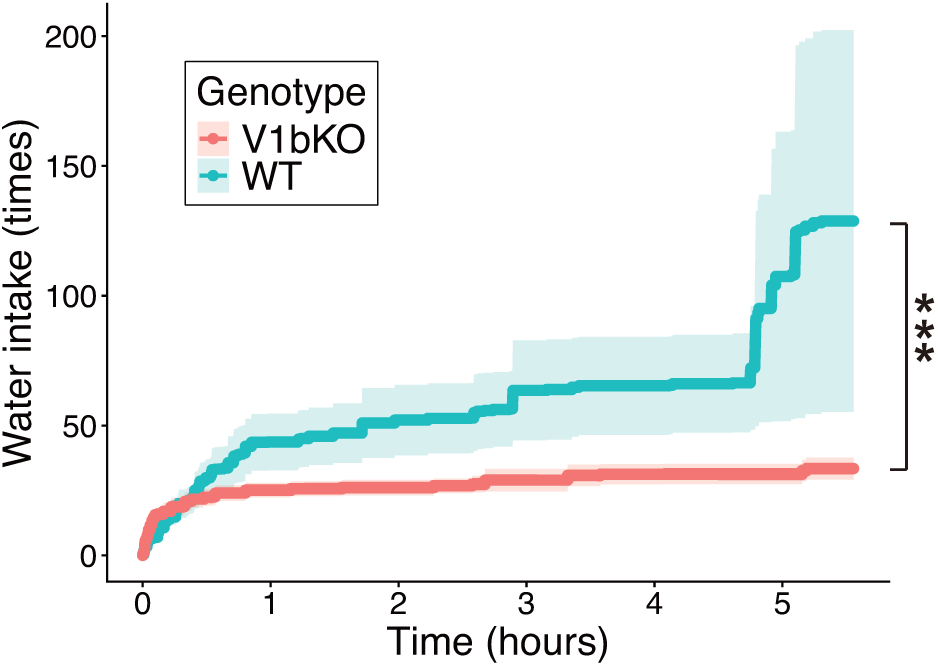
Distinct water access time course in V1bKO mice. Behavioral recordings (6 h) were sampled at 5 fps to quantify drinking patterns in metabolic cages. A deep learning model was used to classify behavior as either ’accessing the water bottle’ or ’away from the water bottle’ for WT (n = 9, green) and V1bKO (n = 9, red) mice. Data are presented as mean *±* s.e.m. of cumulative drinking counts. Asterisks indicate a significant interaction between genotype and time course (***, *p <* 0.001).

We assessed metabolic parameters at the end of the recording period to evaluate physiological homeostasis (Figure 2). Although V1bKO mice began with a significantly lower baseline body weight than WT mice (Figure 2a), percentage body weight reductions were nearly identical between groups at 6 h (V1bKO: 6.1 *±* 1.0%; WT: 6.1 *±* 0.4%) and 24 h (V1bKO: 10.1 *±* 0.9%; WT: 9.5 *±* 2.4%). Two-way ANOVA showed that while both time and genotype significantly affected body weight at 6 h, the effect of genotype was no longer significant by 24 h (Figure 2b, c). Furthermore, when normalized to initial body weight, water consumption and urine excretion were indistinguishable between genotypes at both time points (Figure 2d, e). Blood chemistry parameters, including serum osmolality and sodium levels, remained similar between both groups at baseline (Table 1) and after 6 or 24 h of recording (Figures 2f, g). These findings indicate that V1bKO mice successfully maintain serum osmolality through sufficient water access.

**Figure 2:**
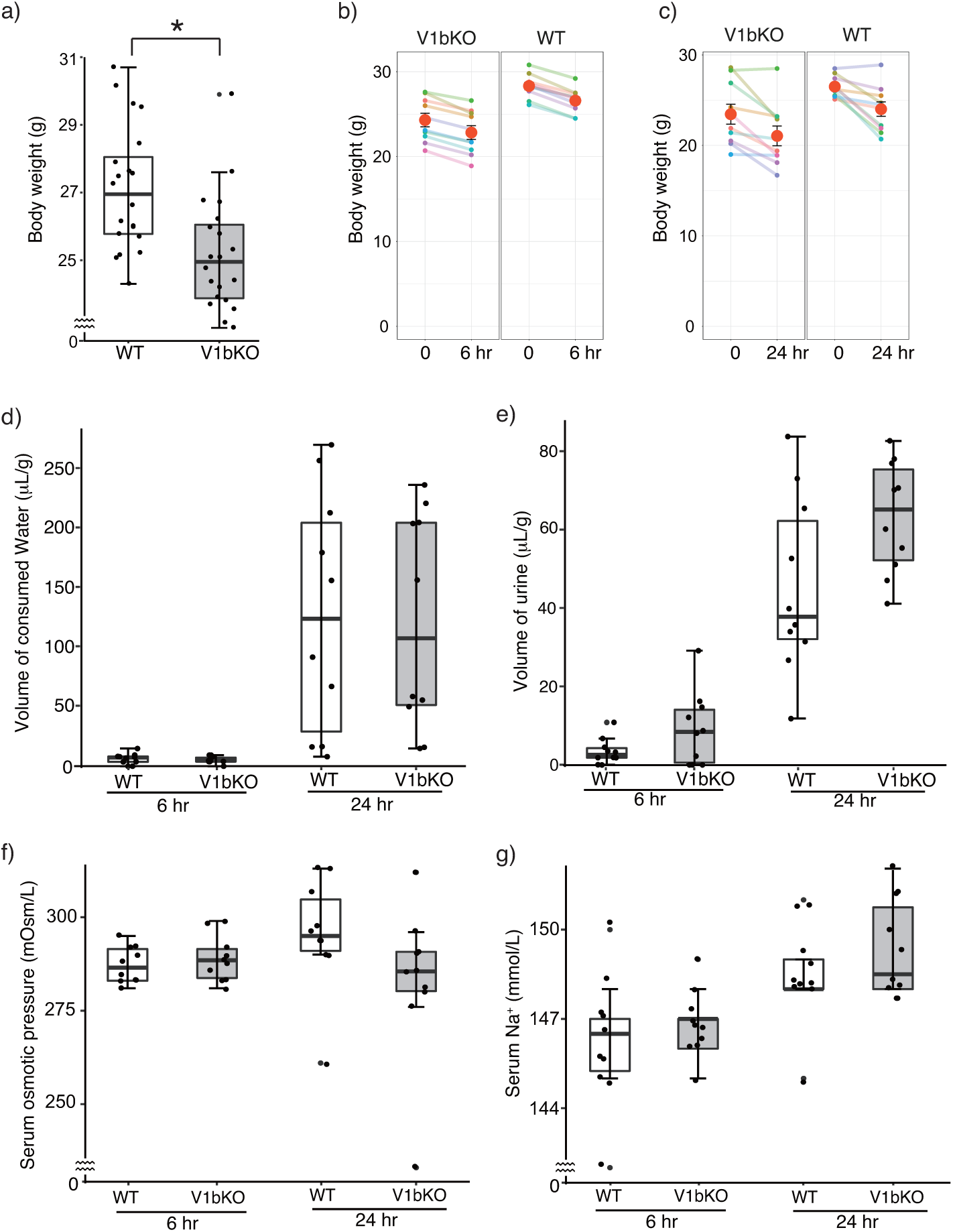
Homeostasis of body water was maintained in the V1bKO mice. (a) Adult V1bKO mice (n = 10) with matched ages showed lower body weights than in the WT mice (n = 10) at baseline. Body weight reduced during the experiments in the metabolic cage for 6 (b) and 24 (c) hours. The mean (red) and s.e.m. are indicated. Volumes of water drunk (d) and urine produced (e), serum osmotic pressure (f) and Na^+^ values in the blood (g) were shown. *, *p <* 0.05

**Table 1:** Serum osmolality and ion concentrations at the start of the experiment.

|  | WT (n = 10) | V1bKO (n = 10) | *p* |
| --- | --- | --- | --- |
| Serum osmolality (mOsm/kg H<sub>2</sub>O) | 323 ± 2 | 324 ± 1 | 0.81 |
| Na<sup>+</sup> (mmol/L) | 150 ± 1 | 149 ± 1 | 0.64 |
| K<sup>+</sup> (mmol/L) | 4.2 ± 0.1 | 4.1 ± 0.1 | 0.60 |
| Cl<sup>-</sup> (mmol/L) | 112 ± 1 | 111 ± 1 | 0.58 |
| Ca<sup>2+</sup> (mmol/L) | 1.19 ± 0.01 | 1.22 ± 0.01 | 0.04* |

\* indicates $p < 0.05$ .

**Table 2:** Blood cell parameters and adjusted p values from post hoc comparisons between WT and V1bKO mice. Values are estimates with standard errors, t-ratios, and BH-adjusted p values (adjusted p). BMT, bone marrow transfer. *, *p <* 0.05 and **, *p <* 0.01.

| Parameters | Basal_BMT | Estimate | s.e.m. | df | t-ratio | p value | adjusted p |
| --- | --- | --- | --- | --- | --- | --- | --- |
| RBC | Basal | -41.8 | 20.5 | 26 | -2.04 | 0.0517 | 0.1 |
| RBC | BMT | -98.6 | 29 | 26 | -3.4 | 0.00218 | 0.009** |
| Hb | Basal | -0.88 | 0.308 | 26 | -2.86 | 0.00826 | 0.03* |
| Hb | BMT | -1.08 | 0.435 | 26 | -2.48 | 0.0199 | 0.03* |
| Ht | Basal | -1.71 | 0.982 | 26 | -1.74 | 0.0933 | 0.2 |
| Ht | BMT | -3.52 | 1.39 | 26 | -2.54 | 0.0176 | 0.03* |
| MCH | Basal | -0.25 | 0.115 | 26 | -2.17 | 0.0391 | 0.10 |
| MCH | BMT | 0.52 | 0.163 | 26 | 3.2 | 0.00364 | 0.009** |
| MCV | Basal | 0.15 | 0.291 | 26 | 0.515 | 0.611 | 0.7 |
| MCV | BMT | 1.14 | 0.412 | 26 | 2.77 | 0.0102 | 0.02* |
| MCHC | Basal | -0.64 | 0.17 | 26 | -3.77 | 0.00086 | 0.007** |
| MCHC | BMT | 0.3 | 0.24 | 26 | 1.25 | 0.223 | 0.3 |
| WBC | Basal | -790 | 867 | 26 | -0.912 | 0.37 | 0.5 |
| WBC | BMT | 260 | 1230 | 26 | 0.212 | 0.834 | 0.8 |
| Plt | Basal | 1.67 | 5.68 | 26 | 0.294 | 0.771 | 0.8 |
| Plt | BMT | 29.3 | 8.04 | 26 | 3.64 | 0.00118 | 0.009** |

### 3.2 Water intake events accumulated more rapidly in V1bKO mice than in WT mice after tolvaptan administration

To investigate the role of the V1b receptor in the water intake response induced by aquaresis, mice were administered the vasopressin V2 receptor antagonist tolvaptan (20 mg/kg). The water intake events increased more rapidly in V1bKO mice than in WT mice (Figure 3a; *β* = 2.52*±*0.85, *z* = 2.98, *p* = 0.0029). The cumulative event curves were fitted using a four-parameter Hill equation to quantify their temporal characteristics (Figure 3b). The fitted pperiods required to reach 50% of the maximal cumulative event count were 2.6 h in V1bKO mice and 3.3 h in WT mice (*p* < 0.05). Thus, the fitted midpoint occurred 0.7 h earlier in V1bKO mice, consistent with a more rapid development of the drinking response. By contrast, the total numbers of water intake events detected during the 6 h observation period did not differ significantly between the groups (V1bKO, 2470 *±* 320; WT, 2191 *±* 407; *p* = 0.59). These findings suggest that deletion of the V1b receptor altered the temporal pattern of the water intake response after tolvaptan administration without significantly affecting the total number of detected water intake events.

**Figure 3:**
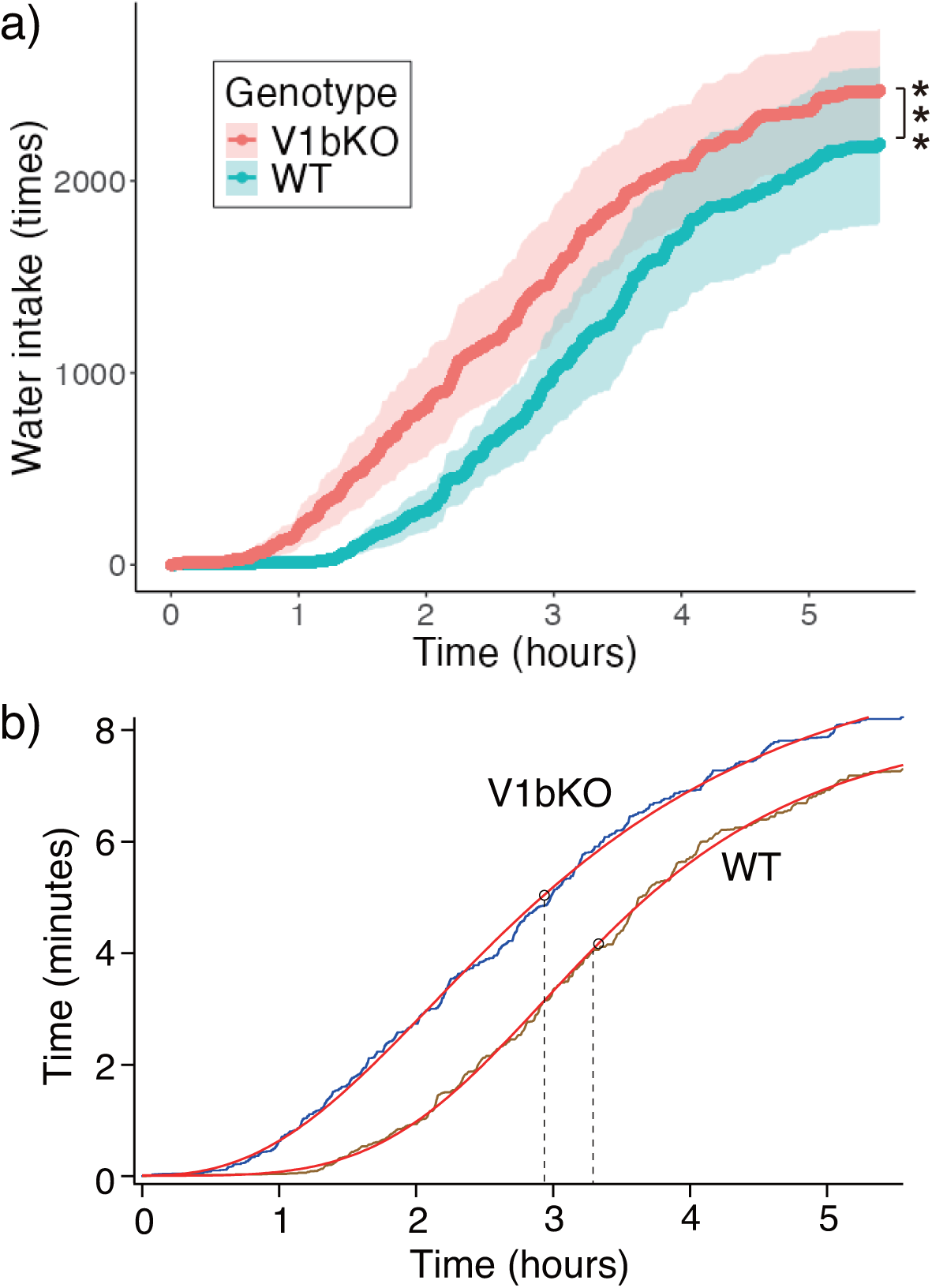
Tolvaptan administration accelerated the frequency of water intake more in V1bKO mice than in WT mice. (a) After tolvaptan (20 mg/kg i.p.), the WT (n = 18) and V1bKO (n = 18) mice were brought individually into the metabolic cage and its behavior was recorded for 6 h. The images were extracted from the video at a rate of 5 images per second; the mice were detected as an object by DNN model. Two types of mouse behavior, water drinking and being away from the water bottle, were classified. The frequency of water drinking was accumulated during the experiments. The genotype of mice had significant influence to the frequency of water intake by GLMM (***, *p* = 0.0029). (b) The accumulating drinking behavior depicted in Figure 3a was fitted to a four-parameter Hill’s equation (red line). The point on the x-axis (crossing with a dashed line) indicates the time point, which gives the half-maximal y-values. In panel (b), y-axis value was recalculated as 5 images per second.

### 3.3 Increased access to the water bottle after tolvaptan administration resulted in the quick recovery of water homeostasis in V1bKO mice

The body water homeostasis was examined 3 h after the tolvaptan administration. Interestingly, the V1bKO mice drank more water during the initial 3 h period and recovered from the reduction of the body weight compared to the WT mice (Figures 4a and 4b). Moreover, serum osmolality and Na^+^ values were reduced compare to those of WT mice, reflecting increased water intake in V1bKO mice. However, urinary volume and osmolality were not different (Figure 4c to 4f). Furthermore, serum Cl^-^ ion concentrations were reduced in V1bKO mice, corresponding to the reduction of Na^+^ levels in V1bKO, while K^+^ levels were the similar in the two groups (Figures 4g and 4h). These results indicate that the Vb1KO mice’s increased access to water reduced high serum osmolality more efficiently than WT mice at this time point.

**Figure 4:**
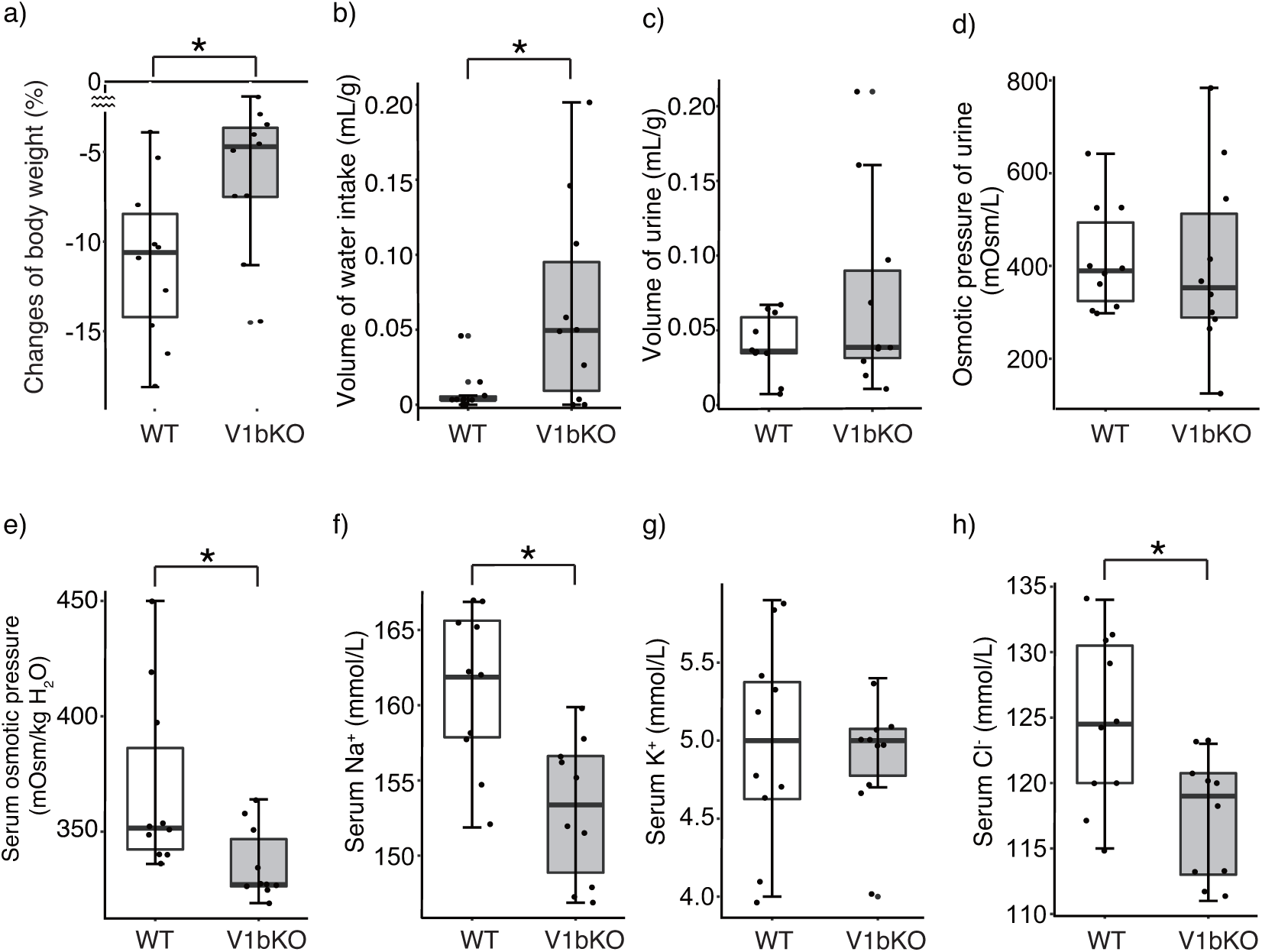
Early initiation of water intake reduced degree of dehydration in the V1bKO mice after aquatic diuresis. Water homeostasis at 3 hours after tolvaptan administration was examined in the V1bKO (n = 10) and WT (n = 10) mice. (a) Changes in body weight, (b) volumes of water consumed, and (c) excreted urine volumes were normalized by initial body weight. Urinary (d) and serum (e) osmolality, and ion levels, (Na^+^ (f), K^+^ (g), and Cl^-^ (h)), were indicated. *, *p <* 0.05

**Supplementary Figure 2:**
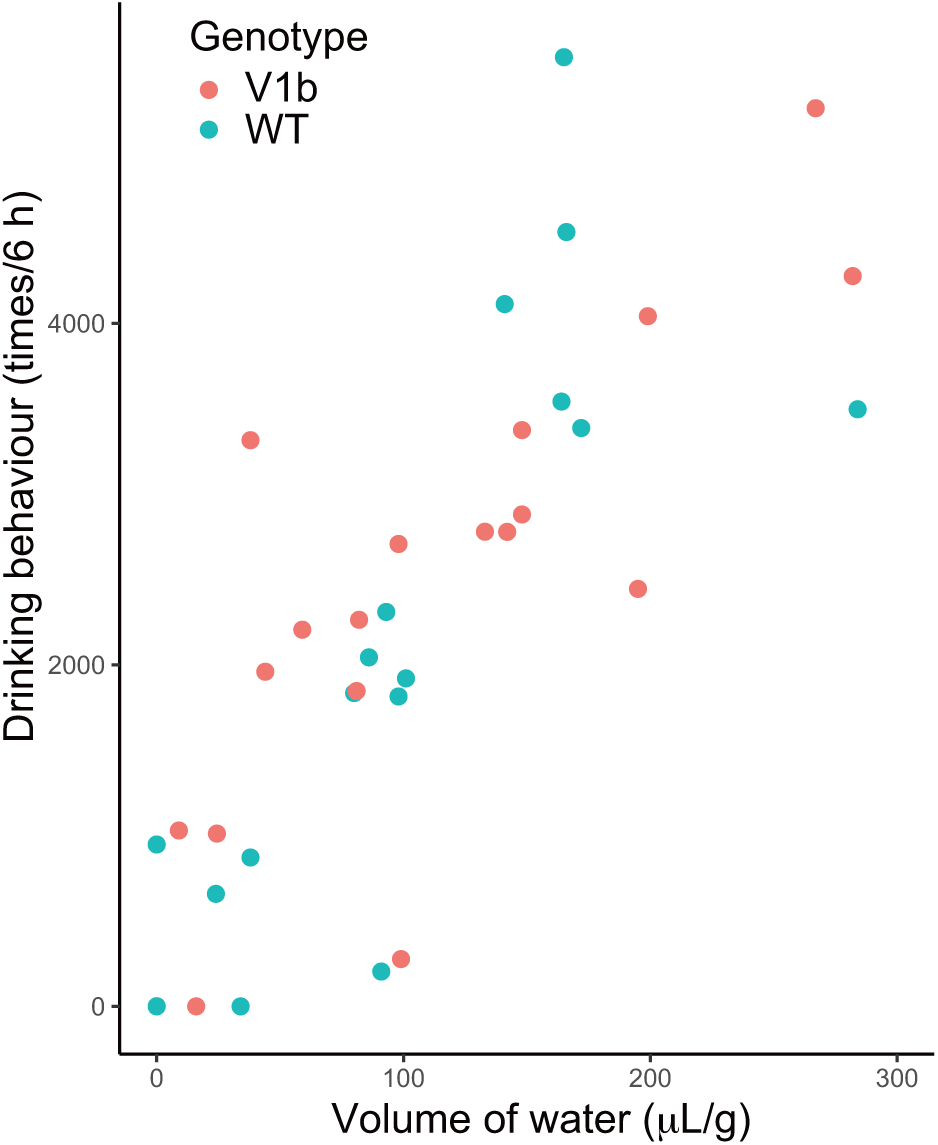
Correlation between the amount of water consumed and the total counts of drinking behavior. X-axis indicates total amount of water consumed and y-axis total counts of access to the water bottle for 6 hours after tolvaptan administration. Points indicate WT (green, n = 10) and V1bKO (red, n = 10) mice.

After an experimental period of 6 h, there was no difference between the two mouse groups in the overall volumes of water consumed (97 *±* 18 and 114 *±* 19 µL/g for WT and V1bKO mice, respectively, n = 18, *p* = 0.49), and excreted urine (109 *±* 16 and 111 *±* 19 µL/g for WT and V1bKO mice, respectively, n = 18, *p* = 0.95). Linear relationship was clearly detected between total counts of drinking behaviors and total amount of water consumed in 6 h in both groups (Supplementary Figure 2).

Because tolvaptan was diluted in 70% ethanol, we administered only ethanol and analyzed water intake behavior for 6 h. After ethanol, both mouse groups showed a similar time course of water intake behavior (Figure 5a). In addition, changes of body weights and volumes of ingested water were of similar levels 3 h after ethanol administration (Figures 5b and 5c). Serum osmotic pressure, but not Na^+^ , K^+^, or Cl^-^ concentrations, increased from basal levels in both groups (Figures 2f and 5e), but was higher in V1bKO than in WT mice (Figures 5d-5g). These findings suggest that ethanol administration did not lead to accelerated drinking, increased water consumption, or improved body weight recovery.

**Figure 5:**
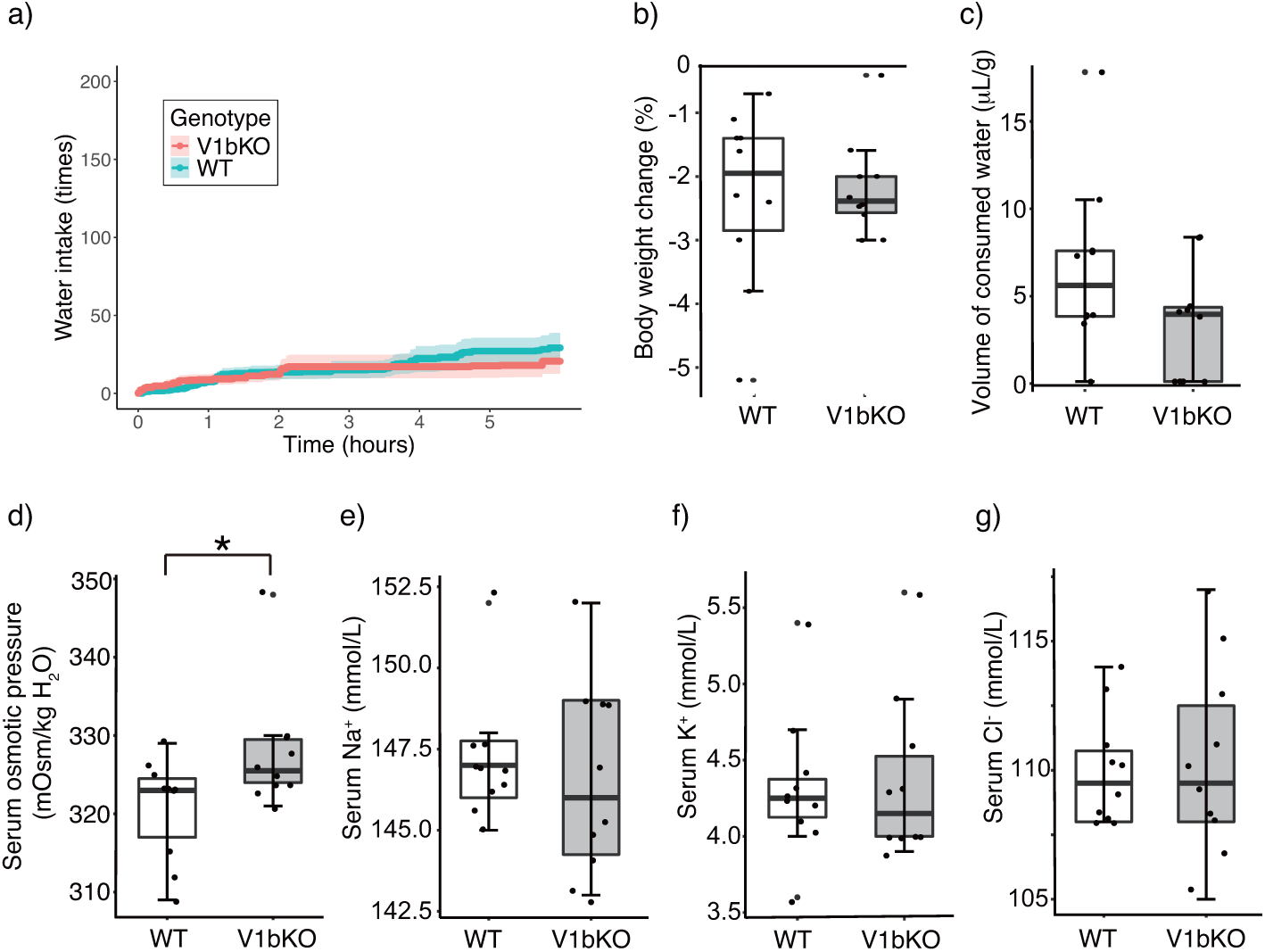
Drinking behavior after ethanol administration was not different between WT and V1bKO mice. After ethanol administration, mice were kept in the metabolic cage for 6 hours for behavior recording (a) and 3 hours for blood and urinary sampling (b – g) from the WT (n = 10) and V1bKO (n = 10) mice under deep anesthesia. *, *p <* 0.05.

### 3.4 Blood Hb level is higher in V1bKO mice than in WT mice

We found that basal Hb and MCHC levels in blood cell counts were significantly higher in the V1bKO mice than in the WT mice (Figure 6b and 6f). Erythrocytosis-like features can be induced by a primary cause originating from BM hematopoietic cells, or by secondary causes, such as increased erythropoietin efficacy or dehydration. Serum erythropoietin was at the same levels in WT and V1bKO mice (279.9 *±* 40.7 and 348.5 *±* 62.1 pg/mL for WT (n = 12) and V1bKO (n = 12) mice, respectively, *p* = 0.37). Because tolvaptan causes strong aquaresis, we examined a possibility that the V1bKO mice’s earlier water intake after tolvaptan treatment might be partly due to a reduced body water reserve compared to WT mice, despite the serum osmotic pressure being the same between the two mouse groups at the basal condition (Table 1) and during observation in the metabolic cage without drug treatment (Figure 2f). Circulating plasma volume was measured by dilution method of Evans blue dye [14]. In preliminary experiments, body weight was correlated with circulating plasma volume; injection of the fixed amount of dye (150 µg) resulted in negative correlation between absorbance of the diluted dye and body weight (Supplementary Figure 3). Therefore, the plasma volume was corrected by body weight of mice. The calculated circulating plasma volume was similar level between WT and V1bKO mice [0.109 *±* 0.009 mL/g body weight and 0.104 *±* 0.010 mL/g body weight for WT (n = 11) and V1bKO (n = 12) mice, respectively, *p* = 0.20].

**Figure 6:**
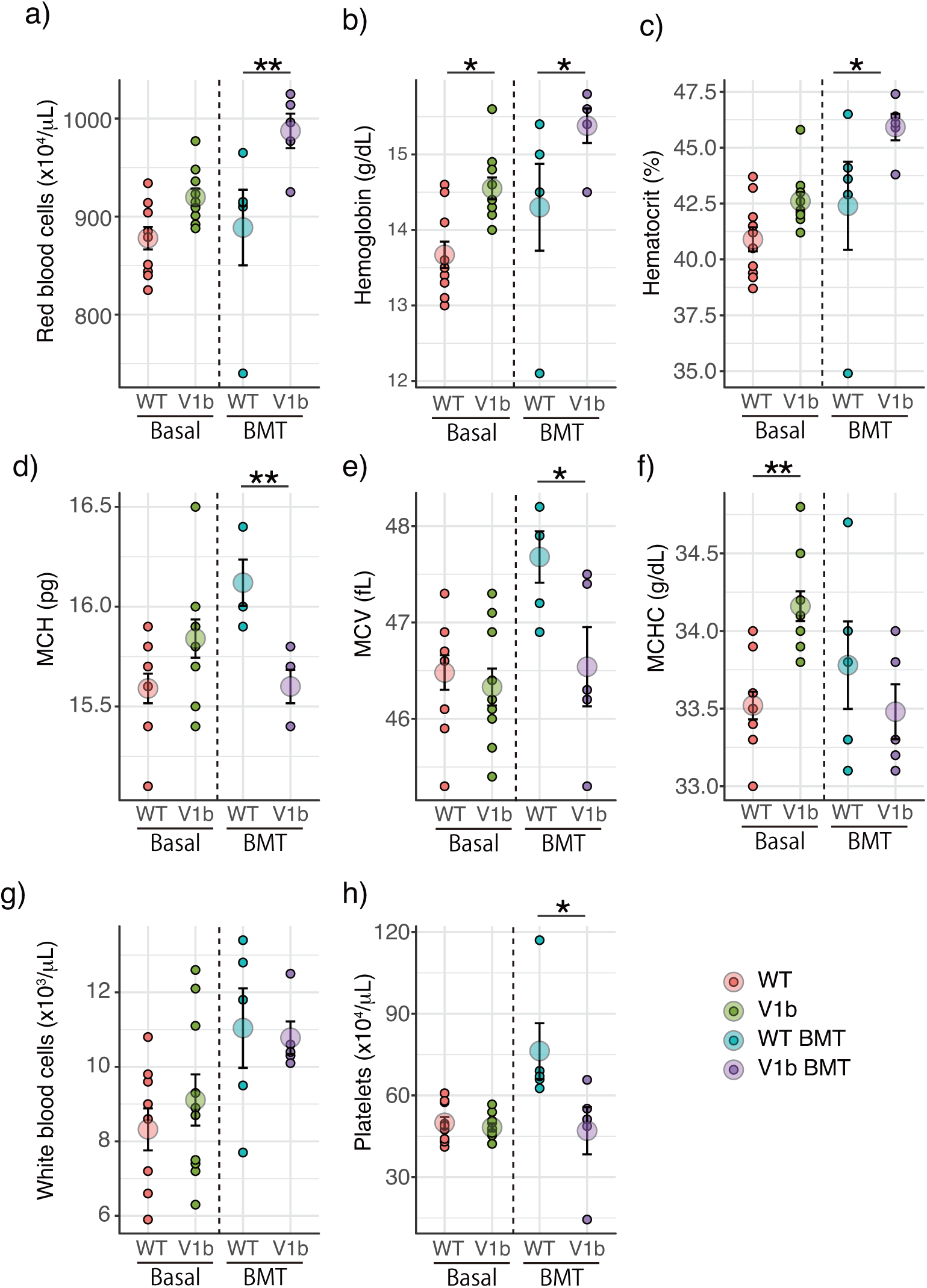
Increased Hb level in V1bKO mice was transferred by transplanting BM of V1bKO mice to irradiated WT mice. Peripheral blood cells were analyzed prior to (left panel, n = 10 for each group) and after (right panel, n = 5 for each group) the bone marrow transplant (BMT) from WT mice (WT BMT) or V1bKO mice (V1b BMT) to irradiated WT mice. The blood sample was obtained 2 months after the BMT. The data shows mean *±* s.e.m. and actual values. *, *p <* 0.05. **, *p <* 0.01.

**Supplementary Figure 3:**
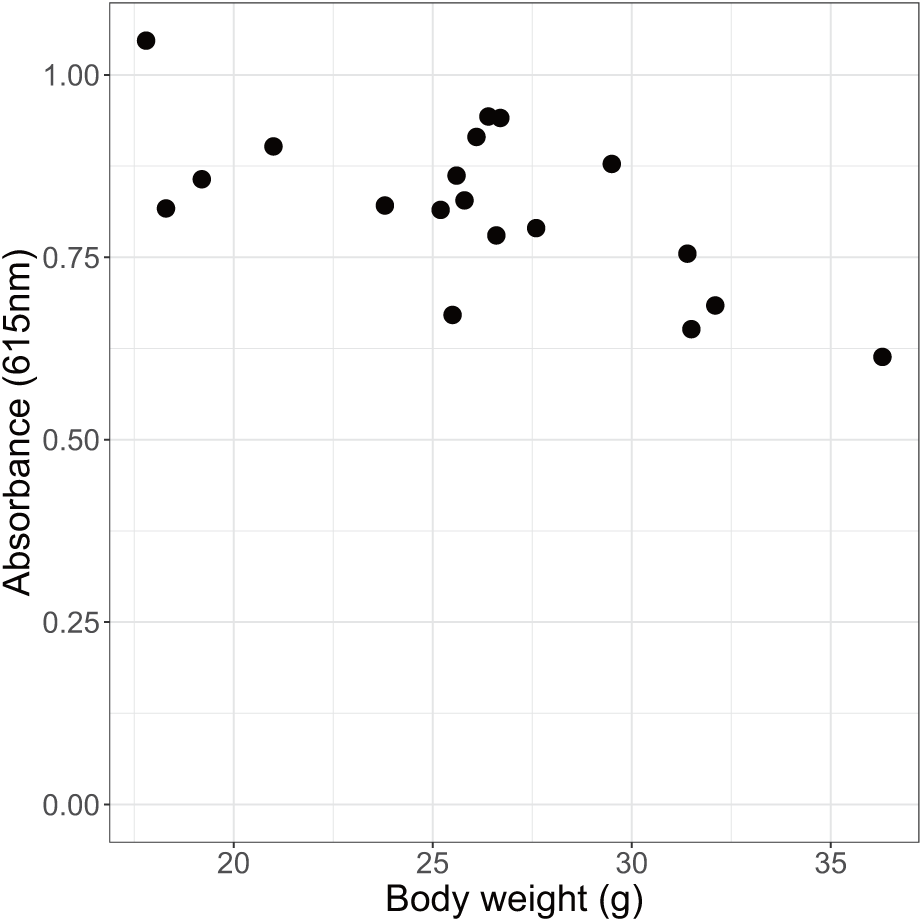
Negative correlation between body weight and absorbance of diluted Evans blue dye in circulating plasma. A fixed amount of 150 µg/25 µL dye was injected into the circulation from right jugular vein of anesthetized mouse. After 10 min, blood was collected from inferior vena cava using heparin-treated syringe and plasma was obtained. The absorbance of the plasma at 615 nm was measured.

To examine the contribution of BM cells to increased Hb in V1bKO mice, the BM of irradiated WT mice was replaced by V1bKO or WT mice’s BM. Two months after the BMT, analysis of the peripheral blood cells revealed an increase in RBC, Hb and Ht levels in WT mice which accepted the BM of the V1bKO mice (Figures 6a–6c). On the other hand, MCH and MCV, but not MCHC, were increased in the mice transplanted with WT BM cells (Figures 6d–6h). Therefore, deletion of V1b receptors in the bone marrow stem cells alter recovery status of transplanted bone marrow and peripheral blood cell populations (Figures 6a and 6c). Based on our results, caution should be exercised when using blood cell parameters to the evaluation of dehydration in V1bKO mice in the current study. Body weights after BMT were at a similar level in mice receiving WT BM (27.6 *±* 1.4, n = 5) and V1bKO BM (28.7 *±* 0.7, n = 5; *p* = 0.12), indicating that the elevated RBC, Hb, and Ht levels in mice with V1bKO BM were not likely due to dehydration or hemoconcentration.

### 3.5 Efficient water intake behavior in V1bKO mice

To evaluate the impact of tolvaptan on V1bKO behavior, we analyzed the spatial dynamics of mouse movement by plotting *xy*-plane trajectories (Figure 7). Representative paths for WT (Figure 7a) and V1bKO (Figure 7b) mice are shown. Overlaid trajectory analysis revealed a reduction in vertical displacement in V1bKO mice compared to WT mice (Figure 7c, d; *n* = 14 for WT, *n* = 16 for V1bKO). This shift was quantified by a significantly lower proportion of vertical movement (Figure 7e) and a compensatory increase in horizontal movement (Figure 7f) in the V1bKO group. Despite these changes in directional proportions, total distance traveled remained indistinguishable between genotypes.

**Figure 7:**
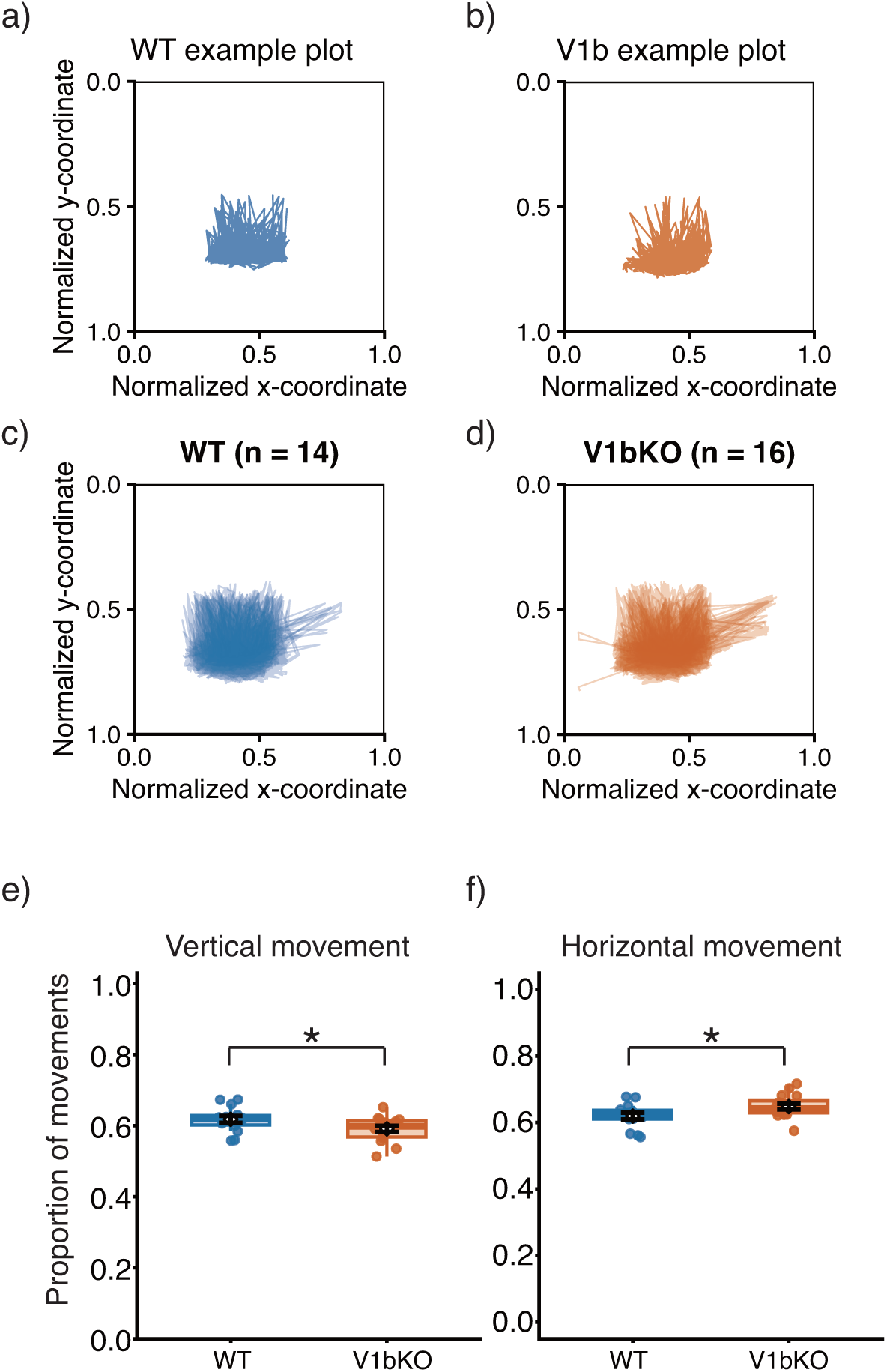
Shift in movement components toward horizontal displacement in V1bKO mice. (a, b) Representative trajectories of single WT (a) and V1bKO (b) mice following tolvaptan administration. (c, d) Group-averaged *xy*-plane traces for WT (*n* = 14, c) and V1bKO (*n* = 16, d) populations. (e, f) Proportion of total movement allocated to vertical (e) and horizontal (f) components. Movements with a relative distance *<* 0.002 were excluded to focus on significant displacement. Error bars represent mean *±* s.e.m. with individual data points overlaid. *, *p <* 0.05.

The biomechanical requirement of upward body extension to reach the drinking nozzle (Supplementary Fig. S1) implies that the diminished vertical displacement in V1bKO mice represents a more efficient drinking posture, as adequate water intake was maintained despite reduced vertical movements. This characteristic exhibited temporal consistency, as the lower proportion of vertical movement was already evident during the first half of the recording sessions (0.39 *±* 0.01 for V1bKO vs. 0.42 *±* 0.01 for WT; *, *p <* 0.05).

## 4 DISCUSSION

In this study, we evaluated the role of V1b receptors in water homeostasis by analyzing metabolic parameters and the temporal patterns of water access in V1bKO mice, under both basal conditions and V2-antagonist-induced aquaresis. In the absence of V2 antagonist treatment, V1b receptor deficiency altered the time course of spontaneous water access (Figure 1) but did not affect the total volume of intake (Figure 2). The efficient drinking behavior observed in V1bKO mice ensured that serum osmolality remained within the same range as that of WT mice after 6 and 24 hours. These findings indicate that V1b receptor function is not essential for the maintenance of osmotic homeostasis or for the regulation of the osmotic satiety set point.

The accelerated accumulation of water intake and sufficient hydration observed in V1bKO mice following tolvaptan administration was an unexpected finding, given that baseline frequencies of spontaneous water access were indistinguishable between genotypes. By utilizing our trained DNN model, we were able to precisely quantify the timing and characteristics of water access. We modeled the relationship between cumulative water intake and time using Hill’s equation, a standard framework for describing dose-response relationships.

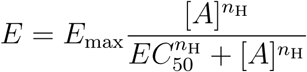

where *E* represents the cumulative number of water access events, *E*_max_ is the maximum observed intake, and [*A*] represents the elapsed time post-administration. The parameter *EC*_50_ denotes the time point at which 50% of *E*_max_ is achieved, and *n*_H_ represents the Hill coefficient, characterizing the steepness of the accumulation curve. Using this mathematical approach, we found that V1b deficiency accelerated the half-maximal response time (EC_50_) by approximately 0.7 hours compared to WT mice under these experimental conditions. This earlier onset of drinking in V1bKO mice was highly effective in mitigating body weight loss and preventing elevations in serum osmolality and electrolyte concentrations, such as Na^+^ and Cl*^−^*. While the Hill’s coefficient traditionally describes cooperative binding (e.g., oxygen-hemoglobin interactions), here it quantifies the degree of sigmoidal cooperativity in the drinking process. We found that the *n*_H_ values for WT and V1bKO mice were significantly different, at 4.0 *±* 0.002 and 2.9 *±* 0.002, respectively. This reduction in *n*_H_ indicates that V1b receptor deletion attenuates the rapid, cooperative-like accumulation of water intake observed in WT mice, despite a faster overall onset of drinking.

Translating these findings to clinical practice could significantly improve the management of water homeostasis during tolvaptan therapy. Thirst is driven by two distinct mechanisms: osmotic thirst, induced by increased plasma osmolality, and hypovolemic thirst, triggered by decreased blood volume [4, 29, 43, 52]. These physiological signals are detected by brain structures such as the SFO and OVLT, which lack a functional blood-brain barrier [39, 38, 22, 23, 35, 36]. Aquaresis is characterized by the loss of free water with relatively minimal sodium excretion [47, 21]. Consequently, tolvaptan can increase serum osmolality, potentially leading to a cascade of diuresis and hypovolemia if water intake does not adequately compensate for urinary losses. Because elderly patients often possess reduced body water reserves and an attenuated thirst sensation [16, 10, 13, 55, 42], they are at heightened risk of dehydration; therefore, careful monitoring and prompt fluid replacement are essential [37, 5, 26, 45]. Our findings suggest that suppressing V1b receptor function during tolvaptan treatment is a safe strategy that may enhance the efficiency of water access and accelerate rehydration.

Previous studies using V1bKO mice have investigated metabolic alterations under both basal conditions and dehydration-induced stress [46, 11]; however, their findings remain inconsistent. For instance, regarding baseline parameters, one study reported increased water consumption and urine excretion in V1bKO mice, whereas another found no significant difference compared to WT mice [46, 11]. Under 24- or 48-hour water deprivation, both studies observed that urinary volumes in V1bKO mice were comparable to those of WT mice [46, 11]. In contrast, the present study demonstrates that during tolvaptan-induced aquaresis, V1bKO mice initiate water intake significantly earlier than WT mice, effectively mitigating the rise in serum osmolality and Na^+^ concentrations. This transient but consistent behavioral shift was uniquely detectable through our high-precision computer vision and deep learning analysis. Indeed, if observations from this study were captured only at a single late time point after tolvaptan administration, the differences in water intake and urine excretion would be undetectable. In addition, the plasma corticosterone level was significantly reduced upon water deprivation in V1bKO mice [46]. The AVP and V1b receptors in the anterior pituitary corticotroph stimulate glucocorticoid secretion in basal and stressed conditions [53]. The balance between the glucocorticoid and mineral corticoid significantly influences body water retention through nuclear receptor activation [17].

The functional interaction between V2 and V1b receptors during AVP-induced dehydration remains a subject of investigation. It has been suggested that renal V1b receptors may modulate V2 receptor-mediated water reabsorption through intracellular signaling crosstalk [25]. In this study, we investigated whether the inhibition of V2 receptors by tolvaptan would reveal a functional role for V1b receptors in regulating water retention. However, following tolvaptan administration, both urine volume and osmolality remained comparable between WT and V1bKO mice. These results indicate that, under the conditions tested, V1b receptor activity did not significantly alter V2-mediated water reabsorption or osmotic balance.

The elevated Hb and MCHC levels observed in V1bKO mice were not attributable to changes in circulating plasma volume or EPO levels (Figure 6). Instead, BMT using V1b- deficient BM cells into recipients resulted in increased Hb, Hct, and RBC counts. These results suggest that the V1b receptor plays a functional role in primary erythropoiesis within the bone marrow. While previous studies have demonstrated that AVP receptors are present in BM cells and that AVP-V1b signaling promotes recovery from anemia [34], our data indicate that V1b deficiency specifically leads to an overproduction of mature RBC parameters (Hb, Hct, and RBC count), suggesting the role of V1b receptor in determining upper limit of mature RBC production. In control WT mice undergoing BMT, the observed increases in MCV and MCH are characteristic of the post-transplant recovery period, driven by the transient release of large, immature reticulocytes [12].

## 5 CONCLUSIONS

This study demonstrates that computer vision analysis, combined with mathematical modeling, provides a high-resolution framework for evaluating the temporal dynamics of drug responses in vivo. We showed that V1b receptor deficiency leads to an accelerated water access following V2 antagonist treatment, potentially enhancing cumulative intake during the early stages of aquaresis. These findings suggest a potential therapeutic strategy for mitigating drug-induced dehydration in humans by modulating the timing of fluid intake. Furthermore, our analysis identified a distinct behavioral signature in V1bKO mice characterized by increased movement efficiency; specifically, these mice achieve necessary physiological tasks̶such as drinking̶with a significantly reduced proportion of vertical movement compared to WT mice. This principle of “efficiency-driven” behavior is consistent with our previous observations, such as the reduced total movement required for maternal pup retrieval in V1bKO mice [50]. Future studies should further investigate the underlying neurobiological mechanisms linking V1b receptor function to these adaptive behavioral patterns.

## Acknowledgements

We would like to thank Ms. Yuki Oyama and Ms. Marie Tanaka for their technical assistances.

## Data Availability Statement

The datasets analyzed in this study can be provided upon reasonable request to the corresponding author.

## Author contribution statement

H.K. and T.K. conceived the project and designed the experiments. H.K., C.S., Y.K. and M.A. performed the experiments. H.K., M.A., F.N., H.T. and T.K. wrote the manuscript. T.K. coordinated and directed the project. All authors analyzed data and reviewed the manuscript.

## Ethics approval statement

Our animal experiments were approved by The Animal Care and Use Committee of the Jichi Medical University and Jichi Medical University Safety Committee for DNA Recombinant Technology. Animal experiments were conducted in accordance with the ARRIVE guidelines (Animal Research: Reporting of In Vivo Experiments).

## Supplementary Data

Supplementary material, including Supplementary Figure 1, 2, and 3 have been provided.

